# Sex Differences in MRI-Based Metrics of Glioma Invasion and Brain Mechanics

**DOI:** 10.1101/2020.11.21.352724

**Authors:** Barrett J. Anderies, Sara F. Yee, Pamela R. Jackson, Cassandra R. Rickertsen, Andrea J. Hawkins-Daarud, Sandra K. Johnston, Kamala R. Clark-Swanson, Joseph M. Hoxworth, Yuan Le, Yuxiang Zhou, Kay M. Pepin, Susan C. Massey, Leland S. Hu, John R. Huston, Kristin R. Swanson

**Author notes:** **Corresponding author:** Barrett J. Anderies, Mayo Clinic Alix School of Medicine.

## Abstract

Gliomas are brain tumors characterized by highly variable growth patterns. Magnetic resonance imaging (MRI) is the cornerstone of glioma diagnosis and management planning. However, glioma features on MRI do not directly correlate with tumor cell distribution. Additionally, there is evidence that glioma tumor characteristics and prognosis are sex-dependent. Magnetic resonance elastography (MRE) is an imaging technique that allows interrogation of tissue stiffness *in-vivo* and has found utility in the imaging of several cancers. We investigate the relationship between MRI features, MRE features, and growth parameters derived from an established mathematical model of glioma proliferation and invasion. Results suggest that both the relationship between tumor volume and tumor stiffness as well as the relationship between the parameters derived from the mathematical model and tumor stiffness are sex-dependent. These findings lend evidence to a growing body of knowledge about the clinical importance of sex in the context of cancer diagnosis, prognosis and treatment.

## Introduction

Gliomas are primary brain tumors characterized by a diverse range of growth patterns and an ability to invade surrounding healthy tissue. Patients diagnosed with glioblastoma (GBM), the highest-grade glioma, have a median survival of only 14.6 months with aggressive standard of care treatment^1^. Magnetic resonance imaging (MRI) is the main imaging modality for visualizing gliomas, including identifying suspected abnormalities, planning targeted treatments, and evaluating treatment response^2^. While MRI provides non-invasive images with excellent soft-tissue contrast, it is non-specific in terms of the full extent of tumor cell invasion. Instead, the abnormalities seen on standard MRI sequences are more reflective of the environmental changes the tumor cells cause rather than the tumor cells themselves^3^. The clinical interpretation of these images (Figure 1) has traditionally been that the primary tumor cell mass is represented by an abnormality on the gadolinium-enhanced T1-weighted (T1Gd) image and that the surrounding T2-weighted-Fluid-Attenuated Inversion Recovery (T2-FLAIR) abnormality is mostly edema with a small amount of invading tumor cells. Practically, this means surgeons target the T1Gd for resection, and radiation and chemotherapy are used to treat the surrounding areas. However, while gliomas are known for their extreme invasion^4^, there is no clinically utilized way to determine an individual tumor’s extent of invasion into the normal appearing brain.

**Figure 1:**
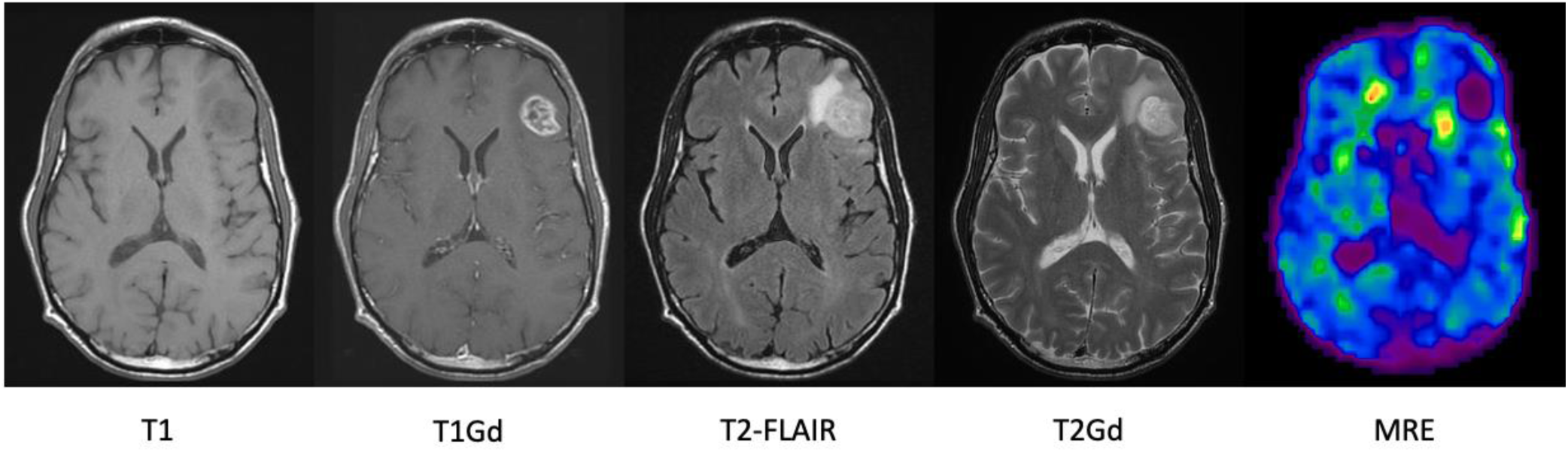
Examples of the different modalities typically used to visualize gliomas for a single patient and an MRE image. In their typical presentation, GBMs are hypointense on T1-weighted (T1) sequences and hyperintense on both T2-weighted (T2) sequences and T2 fluid-attenuated inversion recovery (T2-FLAIR) sequences^2^ all due to a mixture of increase in swelling and extra fluid and additional tumor cells. T1 sequences with gadolinium contrast (T1Gd) show the breakdown of the blood-brain-barrier (contrast leakage) due to the neoplastic process within the tumor, resulting in local hyperintensity.

Swanson et al. developed a personalized imaging-based estimate of tumor invasion using a biomathematical model called the Proliferation-Invasion (PI) model, which returns parameters that describe patient-specific net rates of untreated tumor invasion and proliferation^5–9^. While the model parameters have not been histologically validated due to lack of necessary data, they have been shown to be prognostic of therapeutic response^10,11^, survival^12^, and benefit from extent of resection^13^, as well as predictive of isocitrate dehydrogenase-1 (IDH1) mutation status^14^ and radiation sensitivity^15^. Thus, while not yet used clinically, this metric serves as a surrogate for differentiating more diffuse tumors from more nodular tumors. The prognostic implications of these parameters have also recently been shown to vary based on sex^16–18^. While more studies certainly need to be done, these previous results together with the known biological sex differences in immune system responses support the hypothesis that MRI signal and underlying tumor microenvironmental changes would broadly follow a sex-differentiated pattern^16,19–21^.

Beyond the biological changes tumors cause, they are also known to result in different mechanical properties^22–27^. In the context of the invasive margin of GBMs, it is highly possible that the stiffness of the tissue may be indicative of the different types of abnormalities shown on MRI: swelling due to migrating tumor cells, activated immune cells, extra fluid, and all possible combinations of these phenomena. Magnetic resonance elastography (MRE) was first described in 1995 by Muthupillai et al.^28^ and has emerged as a technique for non-invasively measuring the mechanical properties of tissue. MRE involves inducing shear waves throughout the tissue of interest and measuring the displacement of the tissue within an MRI scanner^22^. MRE has been used to characterize various tissues including liver, skeletal muscle, myocardium, breast and brain^25^. In brain, MRE has found several applications including characterizing the effect age^29^, sex^30^, and dementia^31^ on regional stiffness. In a recent study, Pepin et al. used MRE to demonstrate that gliomas are softer than normal unaffected brain tissue^32^. They further demonstrated that tumors of higher grade were softer than lower grade and that tumors with an IDH1 mutation were stiffer than those with wild-type IDH1. These results are consistent with previous MRE studies of glioma stiffness on MRE^24^, but somewhat surprisingly in the opposite direction of other studies focusing on extracellular matrix stiffness of breast^33,34^ and glioma^23^ tumors. However, the nature of tumor cell invasion in GBMs and the known swelling in the tumor area may help explain these results. This also highlights that the underlying mechanisms connecting cellular biology to gross tumor mechanics and kinetics remain an area of active investigation.

In this paper, we leverage a glioma patient dataset where the patients have received MRE imaging and the PI model to estimated personalized tumor kinetics parameters. We aim to first investigate whether tissue stiffness measured by MRE is indicative of invasive spread and second whether the invasive patterns on MRI/MRE are sex-specific. We first consider how MRI abnormality size, another possible surrogate for invasion, correlates with MRE. We then look at how the imaging-based invasion estimate from the PI model corresponds with MRE values.

## Methods

### Patient Cohort

The patient cohort in this study was previously reported by Pepin et al.^32^ As previously described, preoperative patients suspected of a brain tumor were recruited to the study if they were at least 18 years of age and had an imageable abnormality of at least 2 cm diameter. Enrolled patients had a scheduled date for surgical resection between April 2014 and December 2016. While the original cohort in Pepin et al.^32^ included 18 patients, for this study 10 were excluded due to not having a full set of standard MRIs available (discussed later in methods subsection *Imaging Derived Invasion Metric*, D/ρ). Thus, the included patient cohort consisted of 8 glioma patients (4F,4M) (1 grade II, 3 grade III, and 4 grade IV tumors), 4 of which had IDH1 mutated tumors^32^. Glioma diagnosis, grade and molecular markers (1p/19q codeletion and IDH1-R132H mutations) were determined based on clinical histopathological assessment of surgical biopsies.

### MRI Protocols

We retrospectively analyzed clinical MRI and research MRE images from the MRE glioma patient cohort^32^. Details of the MRI sequence acquisition and region of interest (ROI) generation are outlined in Pepin et al.^32^ and we will briefly review the methods here. The standard anatomic imaging protocol consisted of a T1-weighted inversion recovery echo-spoiled gradient-echo (TR/TE = 6.3/2.8 ms; TI = 400 ms; flip angle = 11°). For the MRE acquisition, a custom passive driver beneath the patient’s head was used to induce shear waves at 60 Hz. During the shear wave motion, the patient was imaged with a spin-echo echo planar imaging (SE-EPI) MRE pulse sequence that synchronized the motion-encoding gradients to the shear waves (TR/TE = 3600/62 ms). Stiffness was computed as previously described^32^. Tissue was assumed to be linear, isotropic, locally homogeneous, and viscoelastic, and the complex shear modulus was computed from the measured displacement fields using 3D direct inversion. The final result was a quantitative map of the tissue shear modulus, from which the sheer stiffness was derived by computing the median magnitude of the complex shear modulus over regions of interest (ROIs).

### MRE Image Segmentation

#### Tumor ROIs

ROIs were manually drawn by an experienced reader using the anatomic imaging sequences (T1, T2, post-contrast T1, etc.) for reference.

#### Contralateral Control (CC)

For each subject, the grouped tumor ROI was reflected to the contralateral hemisphere to identify a personalized control region.

#### MRE

An average brain tissue stiffness value (magnitude of the complex shear modulus (|G*|) in units of kPa) was calculated for each tumor and contralateral control ROI as previously described^32^.

### Imaging Derived Invasion Metric, D/ρ

The MRI-based PI model is a partial differential equation, which quantifies the spatial and temporal growth of tumor cells per unit volume. The model is written mathematically as *c_t_* = ∇ · (*D*∇*c*) + *ρc*(1 – *c/K*) where *c_t_* is the rate of change of tumor cell density in time, *D* is the net rate of invasion (mm^2^/yr), ρ is the net rate of proliferation (/yr), and *K* is the cell carrying capacity of the tissue (cells/mm^3^), which is considered a fixed constant based on an average 10 μm diameter cell. The tumor invasion profile is defined as the ratio of the invasion and proliferation rates, D/ρ. Large values of D/ρ imply diffuse disease, small values of D/ρ imply nodular disease. This model has been used to quantify growth rates for individual GBM patients using MRI data^38,39^.

Standard clinical MRIs (T1Gd, FLAIR/T2) for each subject were segmented to determine their tumor’s diffusion (*D*) and proliferation (*ρ*) values using the PI model^8^. However, 10 patients were excluded from the original MRE cohort published in Pepin et al.^32^ because we did not have access to at least one T2 or FLAIR image and a T1Gd image taken on the same date, which is required for computing D/ρ values. Segmentation was completed using an in-house semi-automated software. Each tumor was measured on T1Gd and T2-FLAIR images, then verified by a second observer. D/ρ was calculated using the PI model^5,8^.

### Statistical Analysis

Student’s t-test was used to test for differences in tumor cell densities based on radiologically defined regions and sex. Pearson’s linear regression was used to analyze correlative relationships between ROI stiffness, CC stiffness, necrotic volume, T1Gd volume, contrast enhancing (CE) volume, FLAIR volume, D/ρ, and patient age. One-way ANOVAs and t-tests were used to compare ROI stiffness, CC stiffness, the difference in tumor and contralateral control stiffness, and D/ρ stratified by sex, tumor grade, and IDH1 mutation status. A p-value of < .05 was considered statistically significant. All calculations were performed in GraphPad PRISM 8 (San Diego, CA).

## Results

### Tumor Size and Stiffness

To investigate the relationship between tumor size and stiffness, we utilized ROIs from the FLAIR image as available, and T2 ROIs were used when they were not (N=2 of 8). Using linear correlation to compare radiographic volumes with measured MRE stiffness values in both the ROI and CC regions, negative trends were observed, though none reached significance (p=0.13 and p=0.07 respectively) (Figure 2). While not statistically significant, R^2^ values were higher for correlations between CC stiffness and both volumes than between ROI stiffness and both volumes.

**Figure 2:**
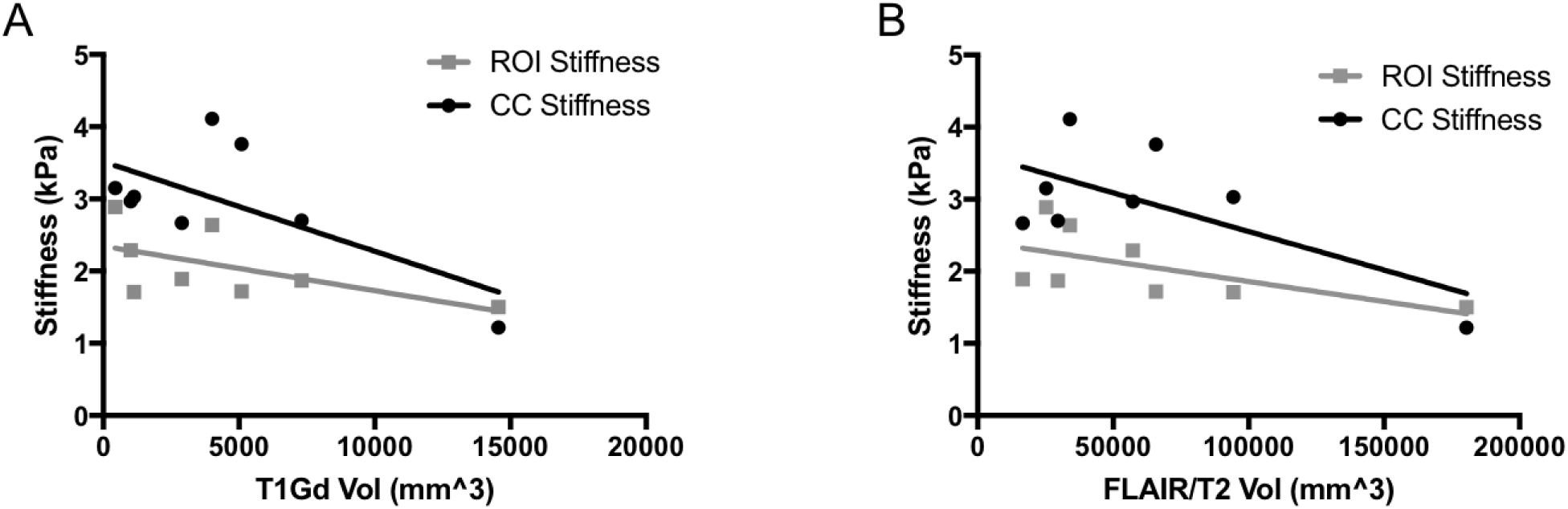
**A**. Linear correlation between T1Gd volume and ROI stiffness (R^2^=0.34, p=0.13) and CC stiffness (R^2^=0.45, p=0.07). **B**. Linear correlation between FLAIR/T2 volumes and ROI stiffness (R^2^=0.37, p=0.11) and CC stiffness (R^2^=0.45, p=0.07).

### Tumor Size and Stiffness Accounting for Sex

Examining this data stratified by sex, separate significant correlations were found between the FLAIR/T2 volumes and ROI stiffness for both sexes (p=0.01 for males, and p<0.01 for females), with females exhibiting a stronger negative slope (−1.7e-5 vs −2.4e-6) (Figure 3). No other relationships were found to be significant.

**Figure 3:**
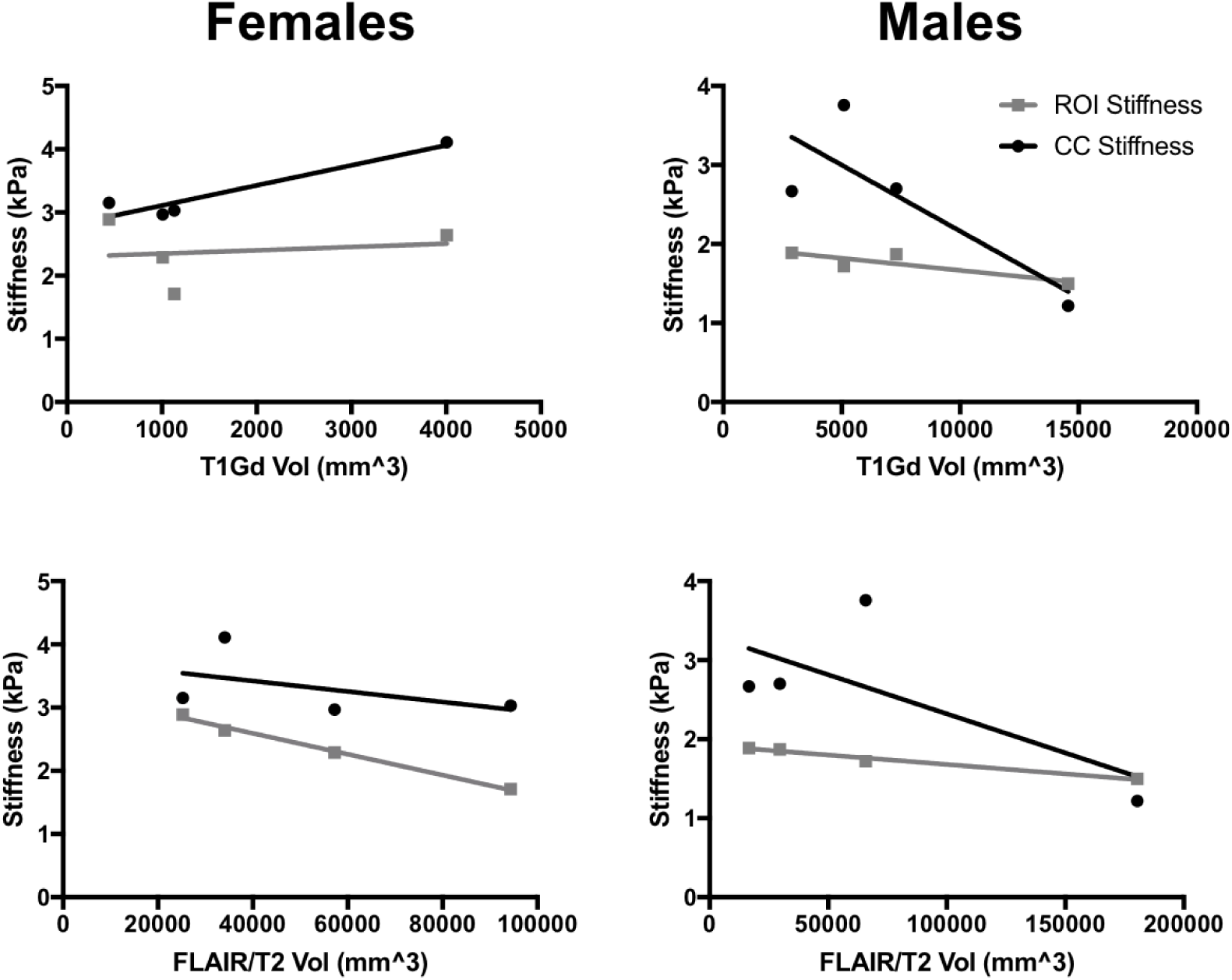
Correlation between stiffness and abnormality volumes considering sex as a variable. Females left column, males right. T1Gd volume top row, FLAIR/T2 bottom. Separate significant correlations seen between the FLAIR/T2 volumes and the stiffness measure. Both sexes had extremely high R^2^ values (0.99 for females and 0.97 for males), but slopes of regression lines were different (−1.7e-5 for females, −2.4e-6 for males).

### D/ρ and stiffness in MRE Patient Cohort

Regression analysis (Figure 4, left) showed no significant relationship between D/ρ and the stiffness calculated in either the CC or ROI (p=0.2673, p=0.2552 respectively). When repeating this analysis in sex-stratified subcohorts (Figure 4, middle), D/ρ did show a significant relationship with stiffness in the ROI region for males (p<0.001) and trended towards significance for females (p=0.057). No significant relationship was found for either sex between D/ρ and stiffness in the CC region (Figure 4, right).

**Figure 4:**
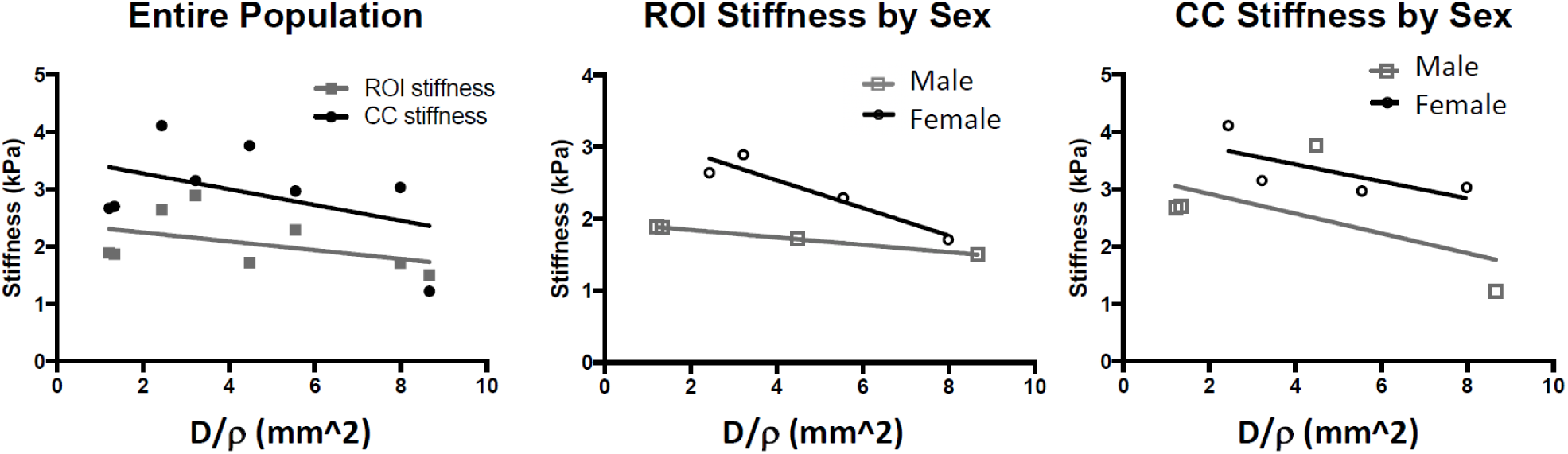
Imaging-based invasion metric and MRE Measured Stiffness. **Left**: Correlation between D/ρ and Stiffness was not significant for either ROI or CC regions (p=0.27 and p=0.25 respectively). **Middle**: Correlation between D/ρ and stiffness in ROI region was significant for males and trending toward significance for females (p<0.001 and p=0.057 respectively). **Right**: Correlation between D/ρ and stiffness in CC region showed no significance for either sex (p=0.42 M, p=0.31 F).

## Discussion

Understanding how the true underlying tumor invasion profile corresponds to radiographic imaging is critical to providing informed clinical care. To date, such studies have been limited due to the difficulty of acquiring tissue for histological analysis. MRI imaging is a proxy measurement of tumors as it primarily resolves contrast (T1Gd) and interstitial fluid accumulation (T2/FLAIR), which are measures of leaky vasculature and edema, respectively. These measures may be correlated with clinical pathology but do not necessarily relate to the local chemical and physical microenvironment within which the tumor cells proliferate and migrate. Pepin et al. previously demonstrated that glioma stiffness decreases with increasing WHO tumor grade and IDH1 mutant gliomas are stiffer than wild-type IDH1 gliomas^32^. In this paper, based on a hypothesis that invasion profiles would influence the tumor stiffness, we have tried to query how MRI signatures on standard pulse sequences correlated with the underlying mechanics and biology.

Our main findings in comparing MRE values with imaging surrogates of invasion, tumor size and an invasion metric based on a biomathematical model, showed that statistically significant correlations were only present when considering patients in sex-specific subcohorts. Specifically, T2/FLAIR volume and MRE in the ROI region was found to be significant when looking at males and females separately but not when pooled. And similarly, the invasion metric, D/ρ, was not significantly correlated with the MRE in the ROI region when all patients were considered together, but it was significant for males alone and was trending towards significance for females. Admittedly, the number of patients we had available for this study due to imaging availability does limit the strength of these results. However, the results suggest that the connection between regions of abnormality as visualized on MRI may relate to the different biological invasion patterns of glioma cells.

Clearly, much research remains to be done. More patients need to be studied and likely the tissue will need to be explored in greater detail to assess cellular composition beyond just the tumor cells. But these results have possible clinical implications. One such area is in assessing drug efficacy. Interstitial pressure is known to negatively influence drug efficacy in solid tumors and Yang et al. determined that standard therapy is more effective in females compared to males^16^. Our results suggest this difference in treatment efficacy may be due to differences in interstitial pressure, particularly the degree and distribution within a given tumor volume. Another implication has to do with the imaging-based invasion metric used here. It is based on a few broad assumptions that the hyperintensity on T1Gd imaging corresponds to regions exhibiting 80% tumor cell density and above while the T2 hyperintense regions correspond to 16% and above (1/5 the T1Gd threshold). The findings we present here suggest that the T2 hyperintense regions may be representative of different phenomena between the sexes. This implies that in the future, the PI model could be more accurate and meaningful if it could be trained in a sex-specific way, using better assumptions of how the imaging regions correlated with tumor cell density.

Clinical imaging remains the primary method of monitoring gliomas and is mostly interpreted with respect to changes in size. Increasing any understanding of how imaging features correspond to tumor characteristics can drastically influence how therapy is chosen and how response to therapy is determined. Much remains to be done, but this work represents a first look at how similar imaging may reflect different tumor invasion depending on the sex of the patient.

## Abbreviations

HGG: high grade gliomas
T1Gd: T1-weighted gadolinium contrasted
T2-FLAIR: T2 weighted fluid attenuated inversion recovery
ROI: region of interest
CC: contralateral control
CE: contrast enhancing
MRI: magnetic resonance imaging;
MRE: magnetic resonance elastography;
MNO: Mathematical Neuro-Oncology Lab;
PNT: Precision Neurotherapeutics Innovation Program
D: tumor diffusiveness
ρ: tumor proliferation
D/ρ: tumor invasiveness

## Acknowledgements

The authors would like to gratefully acknowledge funding for this project through the NCI U01CA220378 and NIH RO1 EB001981. Additionally, we want to acknowledge the work of the Swanson Lab measurement team for generating the ROIs for this paper, with special thanks to Lauren DeGiralamo.

